# Signatures of time interval reproduction in the human electroencephalogram (EEG)

**DOI:** 10.1101/2023.12.11.571138

**Authors:** H. McCone, J.S. Butler, R.G. O’Connell

## Abstract

Accurate timing is essential for coordinating our actions in everyday tasks such as playing music and sport. Although an extensive body of research has examined the human electrophysiological signatures underpinning timing, the specific dynamics of these signals remain unclear. Here, we recorded electroencephalography (EEG) while participants performed a variant of a time interval reproduction task that has previously been administered to macaques, and examined how task performance was predicted by the dynamics of three well known EEG signals: limb-selective motor preparation in the mu/beta band (8-30Hz), the Contingent Negative Variation (CNV) and the Centro-Parietal Positivity (CPP) evidence accumulation signal. In close correspondence with single unit recordings in macaques, contralateral mu/beta signals indicated that participants reproduced intervals by adjusting the starting level and build-up rate of motor preparation to reach a response triggering threshold at the desired time. The CNV showed a highly similar pattern with the exception that its pre-response amplitude was increased for faster reproductions. This pattern of results suggests that, rather than tracing a veridical temporal accumulator as had been suggested in earlier work, the CNV more closely resembles a dynamic anticipatory signal. In contrast, the CPP did not exhibit any relationship with reproduction time suggesting that the evidence accumulation processes guiding perceptual decisions are not involved in generating representations of elapsed time. Our findings highlight close similarities in the dynamics exhibited by intracranial and non-invasive motor preparation signals during interval reproduction while indicating that the CNV traces a functionally distinct process whose precise role remains to be understood.

## 1 Introduction

Accurately judging temporal durations is critical for coordinating our actions in everyday tasks such as hitting an oncoming tennis ball, safely crossing the road, or preparing a meal. A significant body of neurophysiological research has sought to identify the fundamental processes through which the brain tracks the passage of time using tasks in which observers are required to reproduce or discriminate between intervals ranging from hundreds of milliseconds to a few seconds, known as interval timing. Although this research has successfully isolated specific electrophysiological signatures linked to interval timing, there is still a lack of clarity concerning the specific dynamics of these signals.

In non-human primates, behavioural variations in interval timing have been linked to ramping neural activity in cortical areas associated with planning and executing movements (Leon & Shadlen, 2003; Jannsen & Shadlen, 2005). For example, Jazayeri and Shadlen (2015) found that when macaques reproduced interval durations via eye movement responses, a faster build-up rate of spiking in area LIP predicted faster reproduction times. Additionally, LIP activity levels at the start of the reproduction interval were positively correlated with reproduction times. This suggests that in areas involved in preparing the decision-reporting actions, interval timing is achieved via dynamic motor plans that adapt their starting point and build-up rate to reach a fixed threshold at interval offset.

Many human EEG studies of interval timing have focused on a fronto-central negativity known as the Contingent Negative Variation (CNV). Inspired by early models of interval timing, the CNV has been interpreted as a temporal accumulator that indexes a veridical store of elapsed time, with its amplitude representing the amount of time that has passed when reproducing or estimating interval duration (Macar et al., 1999; Casini & Macar, 1999; Macar & Vidal, 2002; Bendixen et al., 2005). However, other studies have failed to find a relationship between CNV amplitude and interval estimates (Elbert et al., 1991; Gibbons & Rammsayer, 2004; Kononowicz & Van Rijn, 2011) and have shown that the build-up rate and peak latency of the CNV is related to interval timing rather than its amplitude (Pfeuty et al., 2005; Macar & Vidal, 2003; Tarantino et al., 2010). Therefore, it remains unclear whether the CNV denotes a veridical accumulator with elapsed time indexed by its amplitude, or if it represents a more flexible build-to-threshold accumulator of the kind highlighted in area LIP.

Beyond the CNV, recent work has highlighted several other candidate signals whose role in interval timing has yet to be investigated. Perceptual decision making research has demonstrated that mu/beta band (8-30Hz) motor preparation signals exhibit many choice-relevant characteristics in common with LIP neurons, including an evidence-dependent ramp to threshold relationship with choice reports and systematic starting level modulations in response to factors such as speed pressure and prior knowledge (O’Connell & Kelly, 2021). In this regard, mu/beta and the CNV appear to be closely linked, as previous mathematical modelling work has highlighted that the CNV’s amplitude at decision onset scales with the starting levels of the decision process and is modulated by experimental manipulations of speed pressure (Boehm et al., 2014). Mu/beta signals would therefore appear likely candidates for tracing the evolution of interval timing decisions but this possibility has yet to be tested.

Another EEG signature of perceptual decision formation that has been extensively studied is the centro-parietal positivity (CPP) which traces the evidence accumulation process independent of the specific sensory or motor requirements of the task (O’Connell et al., 2012; Kelly & O’Connell, 2013). Unlike mu/beta signals, its amplitude at response declines systematically as a function of RT, speed pressure and prior knowledge; while its starting levels remain unchanged (Steinemann et al 2018; Kelly et al 2020). These functional distinctions suggest that the CPP traces a motor-independent representation of cumulative evidence whose amplitude at the time of commitment is determined by strategic influences operating at the motor level (see O’Connell & Kelly, 2021 for full review). The abstract nature of the CPP raises the possibility that it might not be limited to accumulating sensory evidence such as dot motion, but that it could also accumulate temporal pulses. To date, this possibility has not been tested.

The aim of the present study was to determine whether and how the dynamics of the CNV, premotor mu/beta and the CPP predict interval timing. In order to bridge the gap between human and non-human primate work, we used an auditory version of Jazayeri and Shadlen’s (2015) time reproduction task as our behavioural paradigm, and developed our hypotheses with close reference to the motor preparation dynamics those authors observed in macaque LIP. We hypothesised that mu/beta band motor preparation activity would build towards a fixed threshold at response, with starting levels shifted closer to this threshold and a steeper build-up rate for faster reproductions. We hypothesised that the CNV would follow a similar pattern to mu/beta, with a steeper pre-response slope and a larger initial amplitude for faster reproductions. We hypothesised that if the CPP indexes temporal accumulation, it would accumulate towards a fixed bound prior to response, with a steeper build-up rate for shorter reproductions.

## 2 Methods and Materials

### 2.1 Participants

Participants were 25 young adults (12 females, mean age: 25 years, age range: 18-36 years). All participants had normal or corrected to normal vision, had no personal or family history of epilepsy or unexplained fainting, no sensitivity to flickering light and no personal history of neurological or psychiatric illness or brain injury. Two participants were left-handed. Participants provided written consent prior to their participation. The experimental procedures were approved by the Research Ethics Committee of the School of Psychology, Trinity College Dublin in accordance with the Declaration of Helsinki. Participants received €20 for their participation. Participants that were undergraduate psychology students at Trinity College had the option to receive research credits as an alternative to the €20 payment.

### 2.2 Experimental Procedure

Participants sat in a dark, sound attenuated room with their head placed in a chin rest located approximately 45 cm away from a computer monitor. Visual stimuli were displayed at a 1024 x 768 screen resolution and a refresh rate of 60Hz using a 53 centimetre CRT monitor. The task was programmed and presented using MATLAB (Mathworks, Natick, MA) and the Psychtoolbox extension of MATLAB (Kleiner et al., 2007). The sample time intervals presented to participants were 4,800Hz tones, played at a comfortable volume level. Prior to beginning the experiment, participants completed five training blocks. In training blocks, auditory stimuli were presented using Dell AX210 speakers. In experimental blocks, auditory stimuli were presented using Sennheiser HD 650 headphones.

Participants performed a modified version of the Ready-Set-Go time interval reproduction task (Fig 1a) originally developed by Jazayeri and Shadlen (2015). Participants self-initiated each trial by pressing the spacebar. Each trial commenced with the appearance of a centrally presented red dot of 2 pixels on which participants were required to maintain fixation throughout the duration of the trial. Participants were then presented with a sample interval, represented by a single, auditory tone of variable duration (600ms, 650ms, 700ms, 750ms, 800ms, 850ms, 900ms, 950ms or 1000ms with equal probability). Following interval offset, there was a short gap which randomly varied between 350ms and 550ms after which participants were presented with a 100ms cue tone prompting them to reproduce the sample interval by pressing the spacebar such that the time between cue and button press matched their perception of the sample interval. We introduced this gap to create more separation between interval encoding activity and interval reproduction activity due to the potential for signal overlap (e.g. auditory potentials evoked by the offset of the sample tone). Participants performed 16-20 blocks of 45 trials. Each sample interval was presented once every 9 trials in random order.

**Fig 1:**
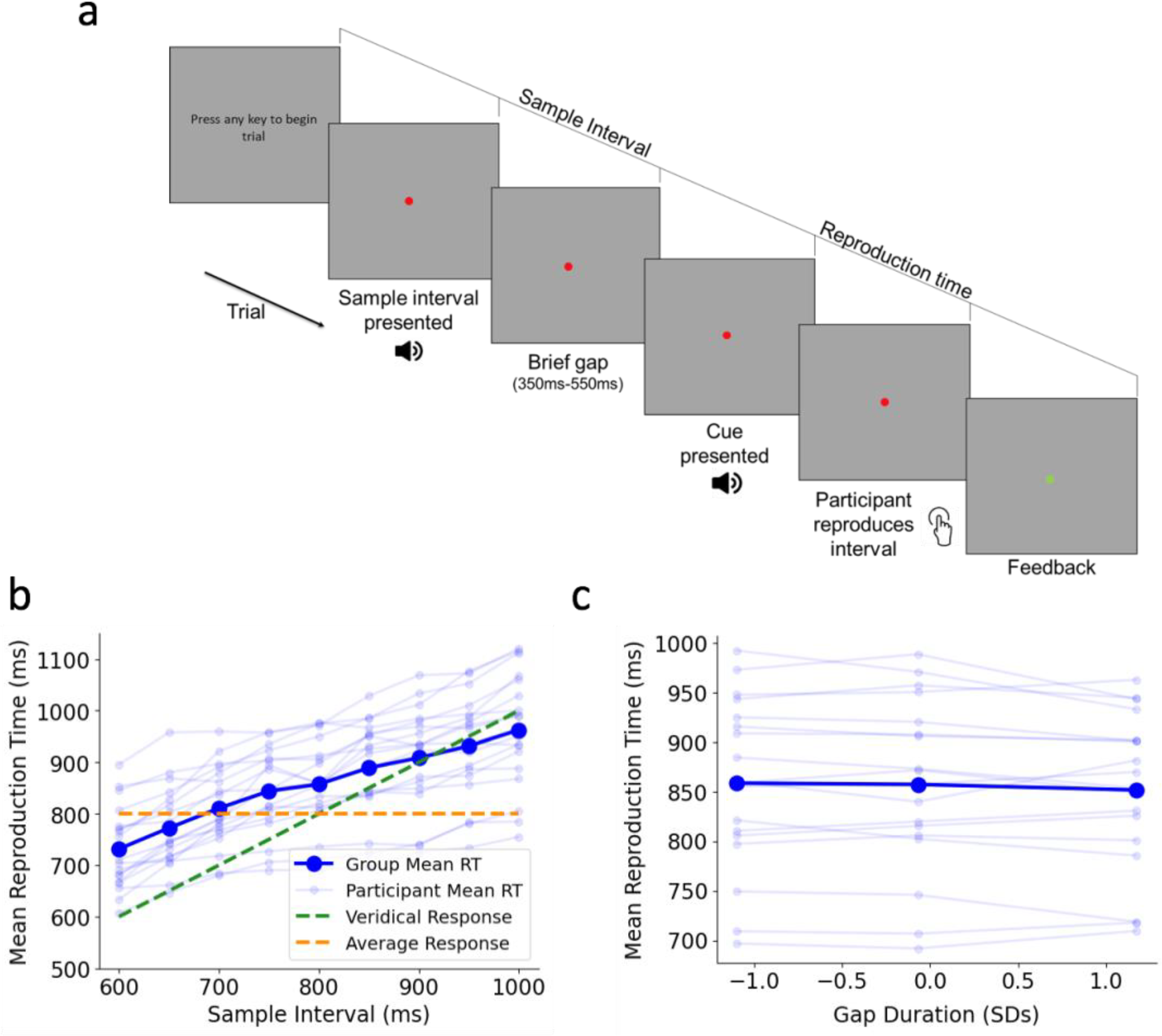
a) Schematic of time reproduction task. Participants were presented with an auditory time interval and reproduced the interval by making a timed keyboard press following a cue. Sample Intervals ranged from 600ms to 1000ms in steps of 50ms. b) Group mean Reproduction Time (dark blue) and individual participant mean Reproduction Time (light blue) as a function of Sample Interval. Reproduction Time increased with Sample Interval, and there was a regression to the mean effect. c) The effect of Gap duration on Reproduction Time. For display purposes, Gaps have been separated into three equal sized bins.

Participants were given positive feedback (fixation dot turned green at the end of the trial) if the reproduced time was within a certain range around the sample interval. To ensure the probability of positive feedback was comparable across sample intervals, the width of the feedback window was scaled with a constant of proportionality, k. The parameter k was initially set to 0.3 at the beginning of the participant’s first training block and was modified throughout the experiment as a function of the participant’s performance via a one-up one-down procedure which subtracted 0.015 from k for each response within this criterion and added 0.015 to k following responses that exceeded it. For trials with responses exceeding this criterion, the fixation dot remained red.

### 2.3 Data Analysis

#### 2.3.1 Behavioural Analysis

We measured participants’ reproduction time (RT) from the midpoint of cue onset (50ms) to the time of response. We removed outlier reproduction times that were less than 150ms and greater than 1500ms (less than 1% of reproduction times). We used a Linear Mixed Model (LMM) to assess the relationship between Sample Interval and Reproduction Time with Sample Interval included as a fixed effect. In addition to fitting random intercepts across participants, the slope of the Sample Interval effect was also included as a random effect. We also included Gap duration in the model as a fixed effect to assess whether it impacted Reproduction Time. Due to a recording error, the distribution of gap durations for five of the participants was shifted by around 100ms (250ms to 450ms). Therefore, we z-scored Gap within-participants so that its value was relative to the distribution of gaps a given participant was presented with. To help the model converge, Sample Interval was z-scored so that Sample Interval and Gap were on the same scale. To obtain standardised beta estimates for the fixed effects, Reproduction Time was also z-scored across participants. Previous studies have established that participants’ reproduction times are biased towards the mean of the underlying distribution of sample intervals. Therefore for each participant, we computed linear regressions with Sample Interval and Gap as predictors, and Reproduction Time as the dependent variable, to examine whether some participants were more veridical responders than others.

#### 2.3.2 EEG Acquisition and Preprocessing

Continuous EEG data were acquired using an ActiveTwo system (BioSemi) from 128 scalp electrodes, digitised at 512 Hz. Data were analysed in custom MATLAB (Mathworks, Natick, MA) and Python 3 scripts (Kluyver et al., 2016), using functions from the EEGLAB (Delorme & Makeig, 2004) and MNE-python (Gramfort et al., 2013) toolboxes. Blinks were detected using two vertical electro-oculogram (EOG) electrodes that were placed above and below the left eye. EEG data were low-pass filtered offline at 30Hz. No offline high-pass filter was applied but EEG data for each block were detrended to remove slow linear drifts. Noisy channels were interpolated and the EEG data were re-referenced offline to the average reference.

EEG data were segmented into three epochs: -200ms to 1100ms relative to sample interval onset, -300ms to 900ms relative to reproduction cue onset, and -1100ms to 100ms relative to the participant’s response time. All epochs were baseline corrected relative to the average activity from -200ms to 0ms preceding sample interval onset. Epochs were rejected if absolute scalp electrode activity exceeded 100μV or bipolar EOG activity (upper minus lower) exceeded 200μV at any time point in the epoch. Six participants were removed from the analyses due to excessive artifacts (fewer than 25 trials per condition), leaving 18 participants for the analyses (9 females, mean age: 24.78 years, age range: 19-36 years). In order to reduce signal overlap, EEG data were transformed into Current Source Density (CSD) following artefact rejection using the CSD toolbox in MATLAB (Kaysar & Tenke, 2006).

#### 2.3.3 Decision Signal Analysis

Based on the grand-average topographies, we measured the CNV from two fronto-central electrodes (located at standard site Fz) centred on the maximum negative grand-average pre-response amplitude. To measure effector-selective Mu/Beta (8-30Hz) activity, we computed a Short-Time Fourier Transform (STFT) with a window size of 375ms (3 full cycles of the lowest mu/beta frequency) and a step size of 25ms. Mu/Beta was measured from four electrodes on the hemisphere contralateral to the hand the participant responded with, centred around the standard sites C3 or C4. As in previous studies examining the CPP, we measured the signal from two central-parietal electrodes, located at the standard site Pz.

We examined the relationship between Sample Interval and the EEG signatures using single-trial LMMs. We focused our analyses of the EEG signatures on three time windows: Sample Interval Offset, Cue Onset, and Reproduction. At Sample Interval Offset, we examined whether the amplitude and build-up rate of the EEG signatures varied with Sample Interval duration. We measured the mean amplitude of the CNV and CPP in a window of -50ms to +50ms from Sample Interval Offset. The build-up rate of each signal was calculated as the slope of a straight line fitted to activity in the same time window. To account for the temporal smearing resulting from the STFT, we measured the mean amplitude and build-up rate of Mu/Beta in a window of -200ms to -100ms before Sample Interval Offset. We assessed the effect of Sample Interval on these measures using LMMs with Sample Interval as a fixed effect, including random intercepts and random slopes across participants as random effects. As in the behavioural LMMs, Sample Interval and the dependent variable were standardized across participants to aid model convergence and obtain standardized beta coefficients for evaluating effect sizes.

Around Cue Onset, we examined whether the amplitude of the EEG signatures scaled with Sample Interval. The amplitude of the CNV and CPP at Cue Onset was measured as the average amplitude in a 100ms window centred at Cue Onset (−50ms to +50ms from Cue Onset). To account for the temporal smearing resulting from the STFT, we measured Cue Onset Mu/Beta at -100ms before Cue Onset. We assessed the effect of Sample Interval on these measures using the same LMM structure as above, with the inclusion of Gap as a fixed effect. Pre-response, we examined whether the build-up rate and amplitude of the signals varied with Sample Interval. The pre-response build-up rate for each of the EEG signatures was quantified as the slope of a straight line fitted to activity from -600ms to -300ms prior to response on each trial. The pre-response amplitude of the CNV and CPP was measured as the average activity between -100ms to -50ms from response. The pre-response amplitude of Mu/Beta was measured as the average activity from -200 to -150ms from response. We assessed the effect of Sample Interval on these measures using the same LMM structure as the Cue Onset analysis.

We also examined the link between these signal measurements and variations in Reproduction Time. Since Sample Interval and Reproduction Time were positively correlated, we median-split the data based on RT (‘Faster’ and ‘Slower’ Bins), with the splits performed within participant and Sample Interval. These models included random intercepts only, as the models would not converge if random slopes were included. As this introduces a risk of overestimating the strength of the fixed effect and underestimating the p-value, we also averaged across trials and performed one way repeated measures ANOVAs with RT Bin as a within-subjects factor.

## 3 Results

### 3.1 Behaviour

As expected, participants adapted their Reproduction Time to the duration of the Sample Interval (Fig1b; x^2^(1) = 31.04, *p* < 0.001, *b* = 0.36), while also exhibiting an overall tendency to overestimate across the entire range of intervals, as well as a regression to the mean effect (Fig1b). The duration of the variable gap between the end of the measurement phase and the beginning of the reproduction phase had a small but significant effect on Reproduction Time (Fig1c: x^2^(1) = 4.99, *p* = 0.025, *b* = - 0.02), with Reproduction Time decreasing as Gap increased (Difference in Group Mean RT between Longest and Shortest Bin in Fig1c = 7ms). We also found that Reproduction Time was not influenced by the preceding sample interval (Fig S1b; x2(1) = 1.34, *p* = 0.25, *b* = -6e^-5^).

### 3.2 Sample Interval Encoding

During Sample Interval presentation, we found no effect of Sample Interval or Reproduction Time on either the amplitude or slope of the CNV, Mu/Beta or CPP measured at the end of the Sample Interval Itself (See left column of Fig2a, b and c; all p>0.05, see S2 for full reporting of statistics).

**Figure 2:**
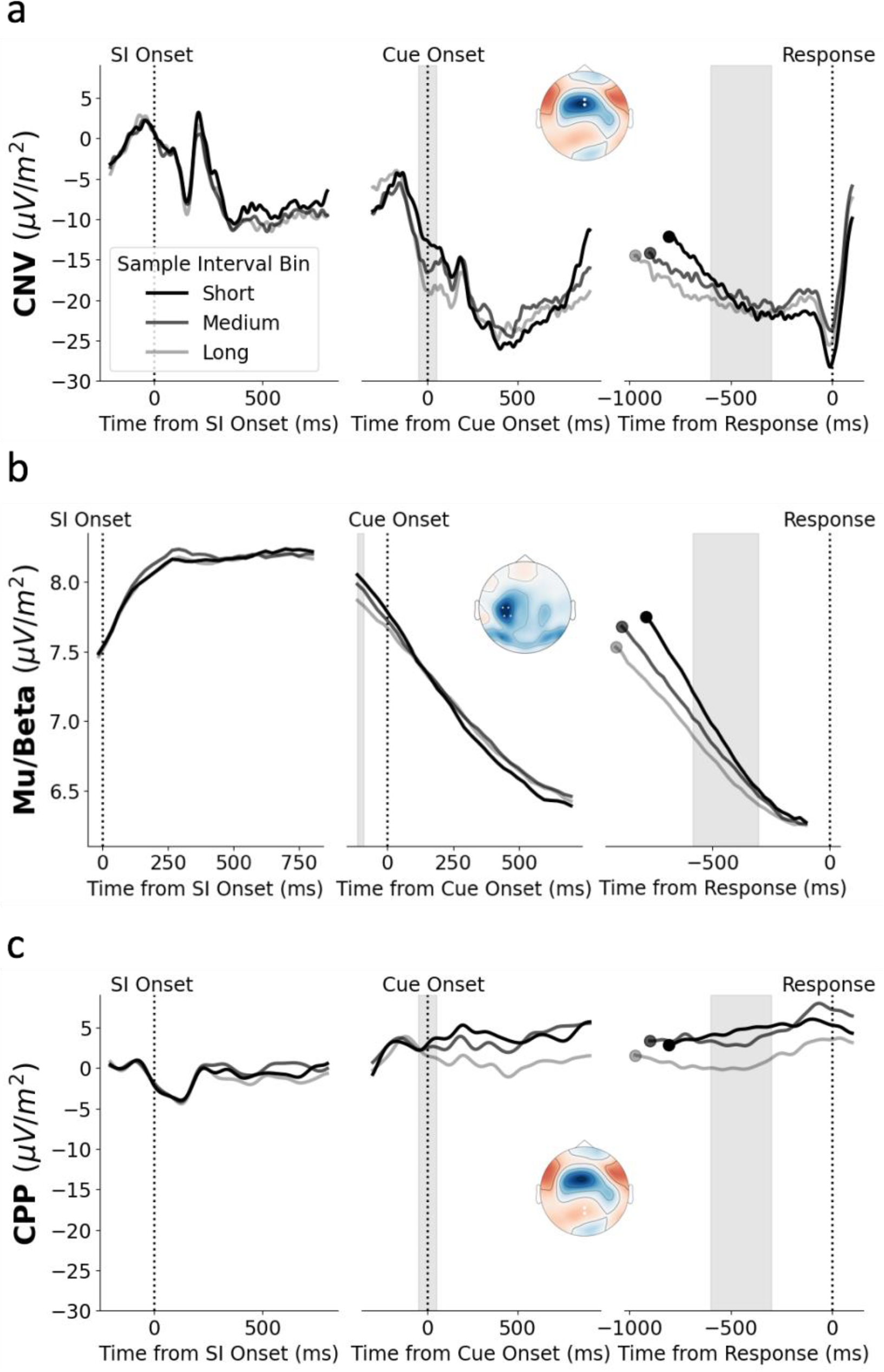
a) CNV as a function of Sample Interval Bin. To aid the visualization of the effect of Sample Interval, we averaged across trials with ‘Short’ (600ms, 650ms, 700ms), ‘Medium’ (750ms, 800ms, 850ms), and ‘Long’ (900ms, 950ms, 1000ms) Sample Intervals. The topography represents the mean grand average activity just prior to the reproduction response (−50ms to 0ms from response). White dots represent the chosen electrodes. Shaded region in the Cue Onset epoch (middle column) represents the time window used for measuring amplitude at cue onset. Shaded region in the response-locked epoch (right column) represents the time window used for measuring pre-response build-up rate. Response-locked waveforms are cropped to the median Cue Onset time relative to the response. b) Mu/Beta as a function of Sample Interval Bin. The topography in this panel represents amplitude at response minus amplitude at Cue Onset. For the purpose of this plot, the electrode positions for left-handed participants were flipped so the left hemisphere in the above topography represents activity that is contralateral to the response hand. c) CPP as a function of Sample Interval Bin. The topography in this panel is the same as in a) apart from the location of the dots representing the chosen electrodes for the CPP.

### 3.3 Reproduction Cue Onset

At Cue Onset, CNV amplitude was larger (more negative) and there was greater Mu/Beta desynchronisation for longer Sample Intervals (Fig2a and 2b; CNV: x^2^(1) = 7.58, *p* = 0.006, *b* = -0.04; Mu/Beta: x^2^(1) = 4.21, *p* = 0.04, *b* = -0.02). While there was no significant effect of Gap duration on CNV amplitude (x^2^(1) = 2.8, *p* = 0.1, *b* = -0.01), Mu/Beta desynchronisation was greater for longer Gap durations (x^2^(1) = 17.81, *p* < 0.001, *b* = -0.02). Additionally, the amplitude of the CNV was larger and Mu/Beta desynchronisation was greater for Faster responses (Fig3a and 3b; CNV: x^2^(1) = 9.51, *p* = 0.002, *b* = -0.03; Mu/Beta: x^2^(1) = 4.75, *p* = 0.03, *b* = -0.01). For these and all following RT Bin analyses, the RM ANOVA result supported the LMM result (see S5 for full reporting of the statistics). This suggests that the starting levels of Mu/Beta and the CNV are adjusted to modify Reproduction Time, with an elevated starting level (more negative CNV and greater Mu/Beta desynchronisation) leading to a faster response. Our analyses also indicated that there was no relationship between CPP amplitude at Cue Onset and either Sample Interval (Fig2c; x^2^(1) = 0.12, *p* = 0.73, *b* = -0.004) or RT Bin (Fig3c; x^2^(1) = 0.44, *p* = 0.5, *b* = 0.006).

**Figure 3:**
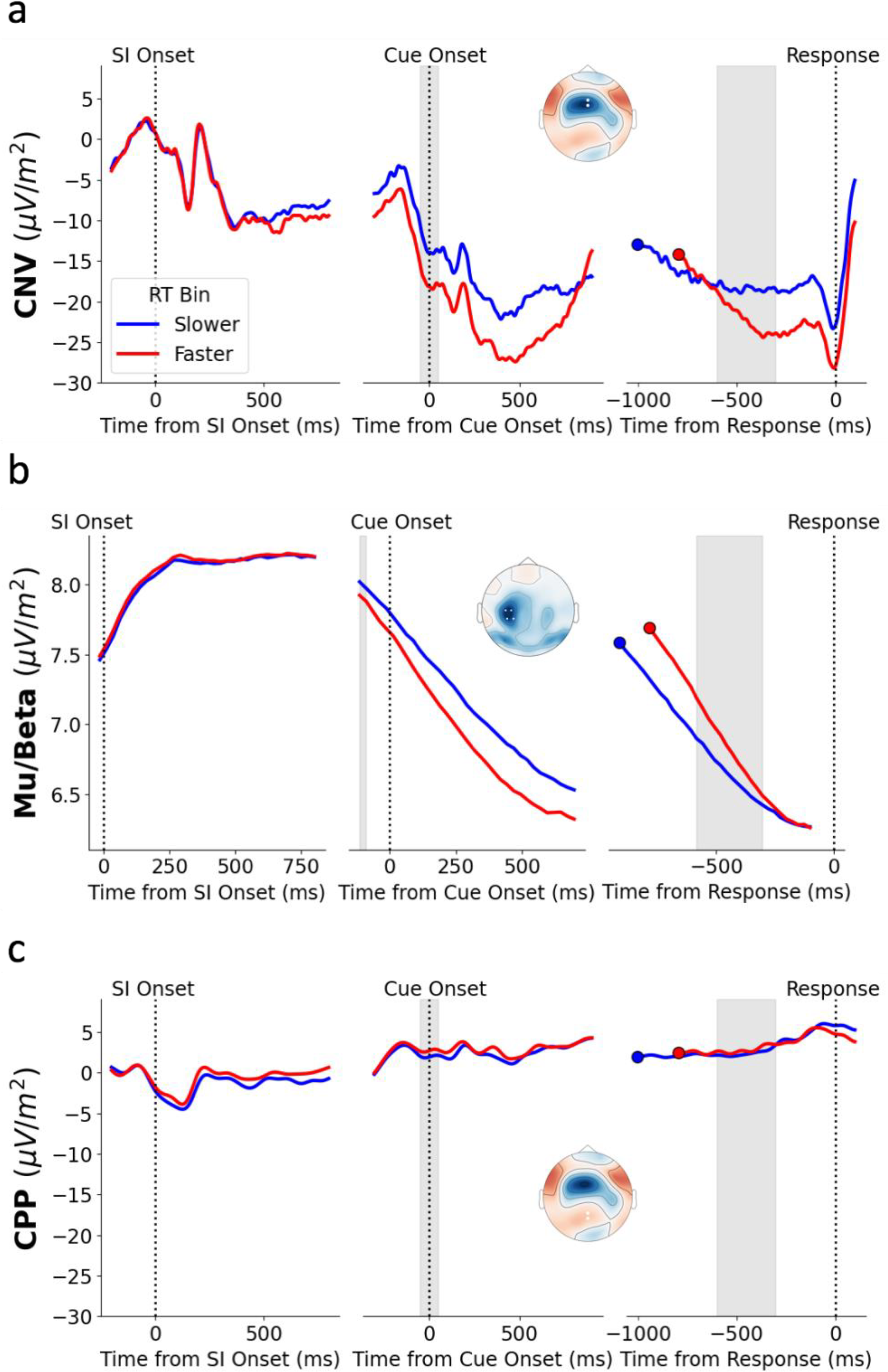
Average waveforms and topographies of the CNV (a), Mu/Beta (b) and CPP (c) across the three epochs as a function of RT Bin. Trials were binned within-participant and Sample Interval into ‘Faster’ (red) and ‘Slower’ (blue) RT Bins based on a Median Split.

### 3.4 Reproduction

Although the pre-response build-up rate of the CNV became shallower as the duration of the Sample Interval increased, this effect did not reach significance (Fig2a; x^2^(1) = 3.1, *p* = 0.08, *b* = 0.03). Similarly, the pre-response amplitude of the CNV did not significantly vary with Sample Interval (Fig2a; x^2^(1) = 0.04, *p* = 0.84, *b* = 0.002). However, there was a significant effect of RT Bin on CNV build-up rate, with a shallower build-up rate for Slower RTs (Fig3a; x^2^(1) = 43.51, *p* < 0.001, *b* = -0.06). Pre-response CNV amplitude was also significantly larger (more negative) for Fast trials (Fig3a; x^2^(1) = 7.67, *p* = 0.006, *b* = -0.02). There was no effect of Gap on CNV build-up rate (x^2^(1) = 2.25, *p* = 0.13, *b* = 0.01) or amplitude (x^2^(1) = 0.001, *p* = 0.97, *b* = 0.0001).

The effect of Sample Interval on pre-response Mu/Beta build-up rate did reach significance (Fig2b; x^2^(1) = 9.64, *p* = 0.002, *b* = 0.05), with Mu/Beta building more slowly for longer Sample Intervals. As with the CNV, the pre-response build-up rate of Mu/Beta was shallower for Slower reproduction times (Fig3b; x^2^(1) = 42.56, *p* < 0.001, *b* = -0.06). Consistent with the idea that Mu/Beta reaches a stereotyped response triggering threshold, pre-response Mu/Beta amplitude did not vary with Sample Interval (Fig2b; x^2^(1) = 1.5, *p* = 0.23, *b* = -0.008) or RT Bin (Fig3b; x^2^(1) = 0.03, *p* = 0.87, *b* = 0.0001). There was a significant effect of Gap on Mu/Beta build-up (x^2^(1) = 8.47, *p* = 0.004, *b* = 0.02), with a shallower build-up rate for longer gaps. Given that longer gaps are associated with a greater Mu/Beta amplitude at Cue Onset, participants may have slightly attenuated their pre-response build-up to avoid responding too early.

Although there was some positive build-up prior to response at central parietal electrodes (Fig2c and 3c), this build-up was attenuated relative to the typically observed levels of pre-response CPP build-up (Kelly & O’Connell, 2013; McGovern et al., 2018). There was no significant relationship between the slope of this pre-response build-up and Sample Interval (Fig2c; x^2^(1) = 0.36, *p* = 0.55, *b* = -0.005) or RT Bin (Fig3c; x^2^(1) = 0.11, *p* = 0.74, *b* = 0.003). These relationships remained non-significant when using a time window closer to the response of (−200ms to -100ms) which is more typically used for measuring CPP build-up rate (Sample Interval: x^2^(1) = 0.11, *p* = 0.75, *b* = 0.003; RT Bin: x^2^(1) = 1.91, *p* = 0.17, *b* = 0.01). Additionally, there was no significant relationship between pre-response CPP amplitude (−100ms to -50ms) and Sample Interval (Fig2c; x^2^(1) = 1.92, *p* = 0.17, *b* = -0.02) or RT Bin (Fig3c; x^2^(1) = 0.03, *p* = 0.85, *b* = 0.002). There was no effect of Gap on CPP build-up rate (x^2^(1) = 1.07, *p* = 0.3, *b* = -0.009) or amplitude (x^2^(1) = 1.16, *p* = 0.28, *b* = -0.009).

As the measurement window for pre-response slope began 600ms prior to the response, some pre-cue activity was included in the slope measurement for trials with reproduction times shorter than 600ms. Therefore, we repeated the above slope analyses for each of the EEG signatures excluding trials with reproduction times shorter than 600ms. Removing these trials from the analyses did not change the above results qualitatively.

## 4 Discussion

The aim of this study was to characterise the human EEG signatures underpinning time reproduction. Specifically, we sought to establish the extent to which the dynamics of the CNV, mu/beta band (8-30Hz) motor preparation activity, and the CPP evidence accumulation signal predict interval reproduction. By using a task that was highly similar to a reproduction task previously used in an investigation of non-human primates, we were also able to determine the degree to which the observed EEG signal dynamics corresponded to those reported in the spiking activity of effector-selective areas that prepare decision-reporting actions. In terms of behaviour, we found that participants adapted their reproduction times to the presented durations. They also exhibited a central tendency effect that is typical of time reproduction. Our EEG analyses suggest that participants reproduced intervals by dynamically adjusting the starting-level and build-up rate of motor preparation so that it reached a fixed response-triggering threshold at the desired time. Specifically, mu/beta band oscillatory activity had starting levels that were shifted closer to threshold and faster build-up rates prior to shorter reproductions. The CNV exhibited a very similar pattern, with larger amplitudes at reproduction cue onset and faster build-up rates for shorter reproduction times. However, the CNV did not reach a fixed threshold at response, as its amplitude was larger for faster reproductions. Unlike the CNV and mu/beta, we found no relationship between the pre-response slope or pre-cue amplitude of the CPP and either interval duration or reproduction time, suggesting that the evidence accumulation processes indexed by the CPP are not involved in interval reproduction.

There is a striking resemblance between the dynamics of the mu/beta motor preparation activity observed here, and the macaque LIP activity identified by Jazayeri and Shadlen (2015). Both signals built towards a fixed threshold at response regardless of sample interval or reproduction time, indicative of a response triggering threshold, and the pre-response build-up rate of both signals varied with reproduction time, with a steeper slope for faster reproductions. Therefore, as suggested by Jazayeri and Shadlen (2015) for LIP, mu/beta in the lead up to response appears to represent a motor plan for reproducing time intervals. Jazayeri and Shadlen (2015) also found that LIP activity around the time of the reproduction cue was increased for longer sample intervals. Here we found a similar effect, as pre-cue mu/beta motor preparation increased with sample interval. In the present study, there was a variable gap between sample interval presentation and interval reproduction which was not included in the Jazayeri and Shadlen (2015) version of the task. We found that motor preparation varied with the duration of this gap, with greater motor preparation for longer gaps. Interestingly, we found the opposite effect when we binned by reproduction time within sample intervals, as pre-cue mu/beta build-up was larger for faster reproductions. This suggests that while there is an increase in pre-cue mu/beta that scales with interval duration, faster reproduction of a given interval is nevertheless associated with greater motor preparation starting levels. This finding is reminiscent of the elevated pre-stimulus mu/beta motor preparation levels associated with faster responding in perceptual decision making tasks (Murphy et al., 2016; Steinemann et al., 2018; Kelly et al., 2020).

In line with more recent electrophysiological studies of interval timing (Kononowicz & Van Rijn, 2011; Ng et al., 2011), the present results suggest that the CNV does not represent a veridical temporal accumulator that accumulates pulses at a fixed rate. Rather than reaching a larger (more negative) amplitude for slower reproduction times as would be predicted by temporal accumulator accounts, the CNV reached a larger amplitude for faster reproductions. The CNV exhibited greater similarities with mu/beta, with starting-levels and pre-response build-up rate that varied with reproduction time. Although the effect did not reach significance, CNV build-up rate also had a similar relationship with sample interval as mu/beta, as it became shallower for longer intervals. These findings suggest that rather than representing a veridical accumulator signal, the CNV more closely resembles a dynamic anticipatory signal like the urgency signals identified in perceptual decision making research (Hanks et al., 2014; Murphy et al., 2016). Indeed, the pre-cue effect of a larger CNV amplitude for faster reproductions closely resembles the larger pre-evidence CNV amplitude observed under increased speed pressure in perceptual decision making that has been linked to reduced response caution (Boehm et al., 2014). In both cases, reduced time to respond is associated with a larger CNV prior to the response window, suggesting that the process reflected in the CNV plays a role in speeding up or slowing down responding. However, the CNV did not reach a stereotyped threshold at response, suggesting that it is not a build-to-threshold motor planning signal like mu/beta and LIP. Additionally, the CNV is implicated in perceptual timing tasks such as the bisection task, in which participants categorise time intervals without making a motor response (see Kononowicz & Penney, 2016; Van Rijn et al., 2011 for reviews). Therefore, it seems that the CNV is not solely linked to the planning and executing of motor responses, but is more closely aligned with the processes involved in cognitive deliberation.

Rather than building until the reproduction response was made, the CNV plateaued around 250ms prior to the response and exhibited a return toward baseline around 100ms prior to the response. Although it did not reverse its trajectory like the CNV, mu/beta also exhibited a deceleration in build-up rate prior to the response. Further work will be required to probe these dynamics and the functional distinctions between the CNV and mu/beta. One speculation that could be examined is that the amplitude of the CNV indexes an urgency process that specifies the rate at which motor preparation should increase.

Given that the CPP has been shown to index the accumulation of sensory evidence in both visual and auditory modalities across a range of perceptual decision making tasks, we hypothesised that it may also represent a temporal accumulator signal. However, the present results suggest that the CPP does not index the accumulation of temporal pulses, as neither its slope or amplitude were related to sample interval or reproduction time. One possibility is that the CPP is restricted to accumulating samples of sensory evidence when the decision maker has to choose between multiple alternatives. Alternatively, the CPP may only accumulate samples of evidence when an external stimulus is presented, such as an auditory tone. In the absence of external sensory input, as in the reproduction phase of our task, the CPP may not accumulate an internal signal representing elapsed time. Future work could address this question by utilising the same task we used here, but with the addition of an auditory tone played while participants reproduced the interval.

Whereas Jazayeri and Shadlen (2015) found that LIP activity continued to build throughout the sample interval at a decelerating rate, here motor preparation and the CNV appeared to plateau midway through the sample interval. Additionally, we did not find a relationship between interval duration and the build-up rate or amplitude of any of the three signals at sample interval offset. A possible explanation for this apparent discrepancy lies in our adding of a variable gap between sample interval offset and reproduction onset. The fact that observers in Jazayeri and Shadlen (2015) could begin reproduction as soon as the sample interval ended meant that they could continually prepare and tailor their motor plan during the sample interval itself. Here, the variable gap would ensure that such a strategy would not be as beneficial and our data suggest that participants waited until the end of the sample interval to begin preparing their responses. The fact that mu/beta and the CNV did exhibit amplitude modulations immediately prior to cue onset that scaled with sample interval and reproduction time suggests that some anticipatory preparation did occur, but over a narrower interval.

In conclusion, our results extend previous work conducted in macaques to human participants, showing that humans reproduce time intervals by flexibly adjusting motor preparation activity to hit a response triggering threshold at the desired time. Furthermore, our findings support a growing body of literature indicating that the CNV does not represent a veridical temporal accumulator. Instead, the CNV bears a striking resemblance to motor preparation, suggesting that it plays a role in speeding up and slowing down responding. Our results also indicate that the evidence accumulation processes that govern perceptual decision making do not play a role in interval timing.

## Data and Code Availability

Preprocessed EEG data and analysis code are freely available online.

## Author Contributions

McCone, H., Butler, J. S., and O’Connell, R. G. designed the study, analysed the data and prepared the manuscript. McCone, H. collected the data.

## Funding

This research was funded by a grant from the European Research Council (ERC): European Research Council Consolidator Grant IndDecision – 865474

## Declaration of Competing Interests

No conflicts of interest.

## Acknowledgements

Thank you to Pat McKeown and Daria Monakhovych for assisting with data collection. Thank you to all the participants who kindly participated in the study.

## Notes

https://osf.io/3vtg9/?view_only=300d5c1e3d7e4891a9ba19e209fe5ccd

### Competing Interest Statement

The authors have declared no competing interest.

