## Supplementary Materials for "Signatures of time interval reproduction in the human electroencephalogram (EEG)"

#### S1: Strength of sample interval effect varied across participants; previous trial sample interval did not predict reproduction time.

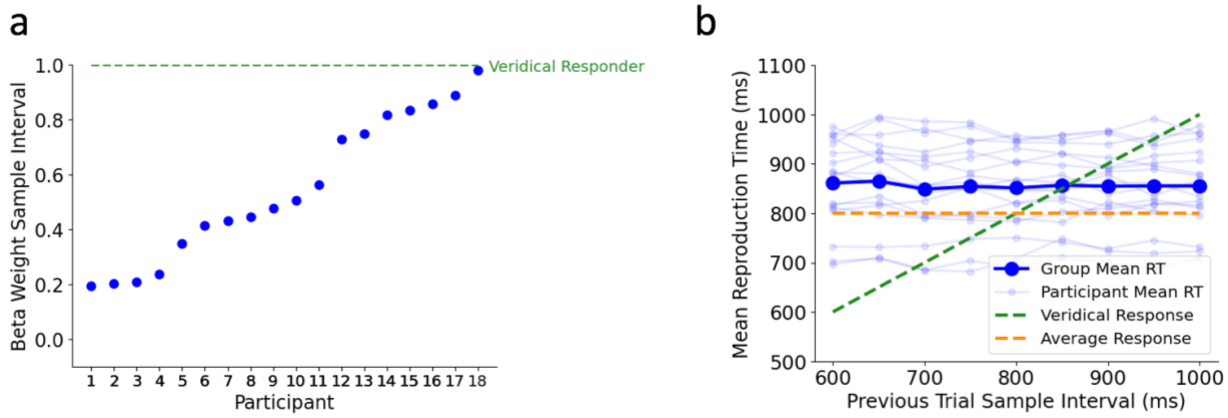


Figure S1: a) The degree to which Reproduction Time was influenced by Sample Interval varied substantially across participants. Each point represents the beta coefficient for Sample Interval in a single participant linear regression predicting Reproduction Time. b) Group mean Reproduction Time (dark blue) and individual participant mean Reproduction Time (light blue) as a function of previous trial sample interval duration.

#### S2: EEG Signatures at Sample Interval Offset

- No effect of Sample Interval on CNV amplitude ($x^2(1) = 0.12$, $p = 0.73$, $b = 0.005$) or slope ($x^2(1) = 0.21$, $p = 0.65$, $b = 0.005$).
- No effect of RT Bin on CNV amplitude ($x^2(1) = 2.71$, $p = 0.1$, $b = -0.006$) or CNV slope ($x^2(1) = 0.005$, $p = 0.94$, $b = 6.28e-4$).

- No effect of Sample Interval on Mu/Beta amplitude ($x^2(1) = 0.006$, $p = 0.94$, $b = -8.5e-4$) or slope ($x^2(1) = 0.77$, $p = 0.38$, $b = -0.01$).
- No effect of RT Bin on Mu/Beta amplitude ($x^2(1) = 8.23e-7$, $p = 0.97$, $b = -1.47e-4$) or slope ($x^2(1) = 0.034$, $p = 0.85$, $b = -0.002$)

- No effect of Sample Interval on CPP amplitude ($x^2(1) = 0.19$, $p = 0.66$, $b = 0.005$) or slope ($x^2(1) = 0.11$, $p = 0.74$, $b =-0.003$).
- No effect of RT Bin on CPP amplitude ($x^2(1) = 0.44$, $p = 0.51$, $b = -0.002$) or CPP slope ($x^2(1) = 5.86e-4$, $p = 0.98$, $b = -7.46e-4$).

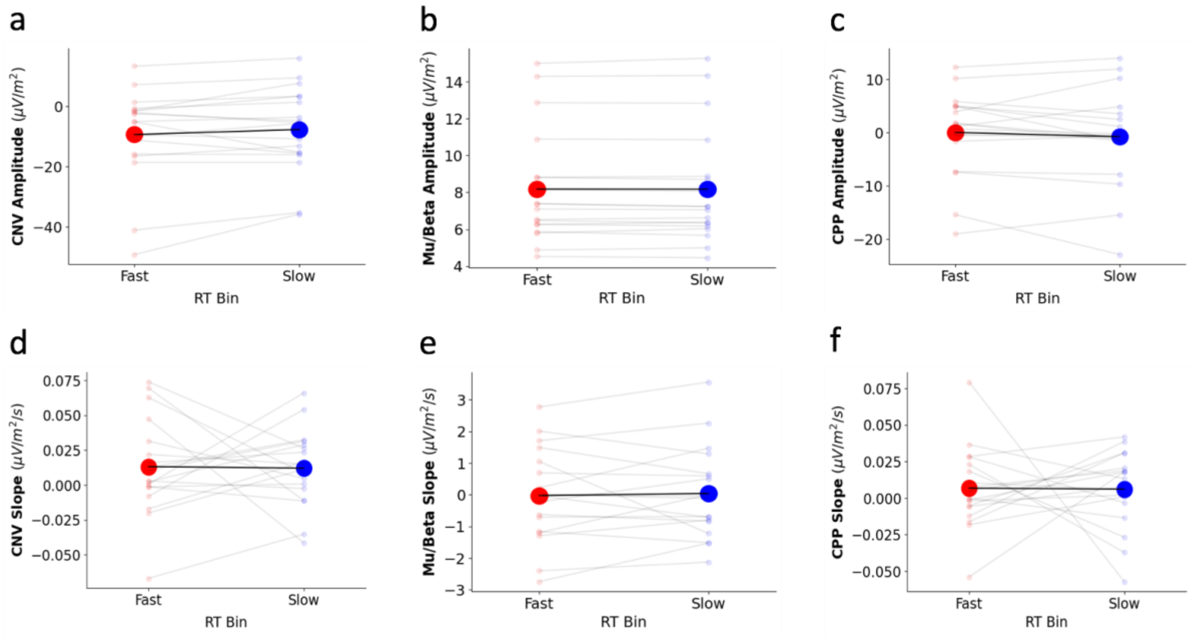


Figure S2: Group mean (black lines and large markers) and participant mean (grey lines and small markers) amplitude (first row) and build-up rate (second row) as a function of RT Bin at sample interval offset. First column is CNV (a and d), second column is Mu/Beta (b and e) and third column is CPP (c and f).

#### S3: Waveforms Separated by Sample Interval

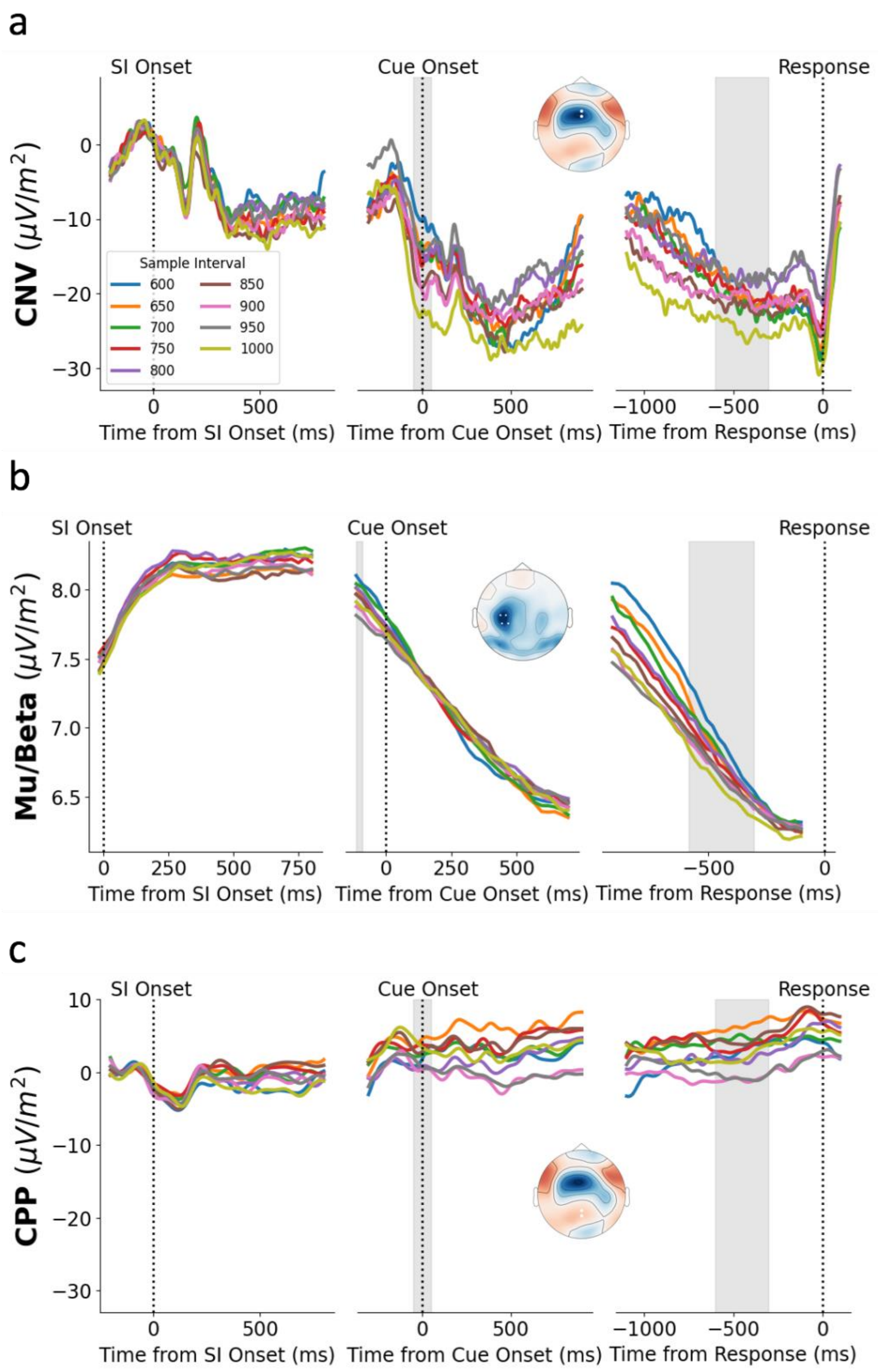


Figure S3: Average waveforms and topographies of the CNV (a), Mu/Beta (b) and CPP (c) across the three epochs as a function of Sample Interval.

### S4: Cue Onset and Pre-Response Measure Lineplots

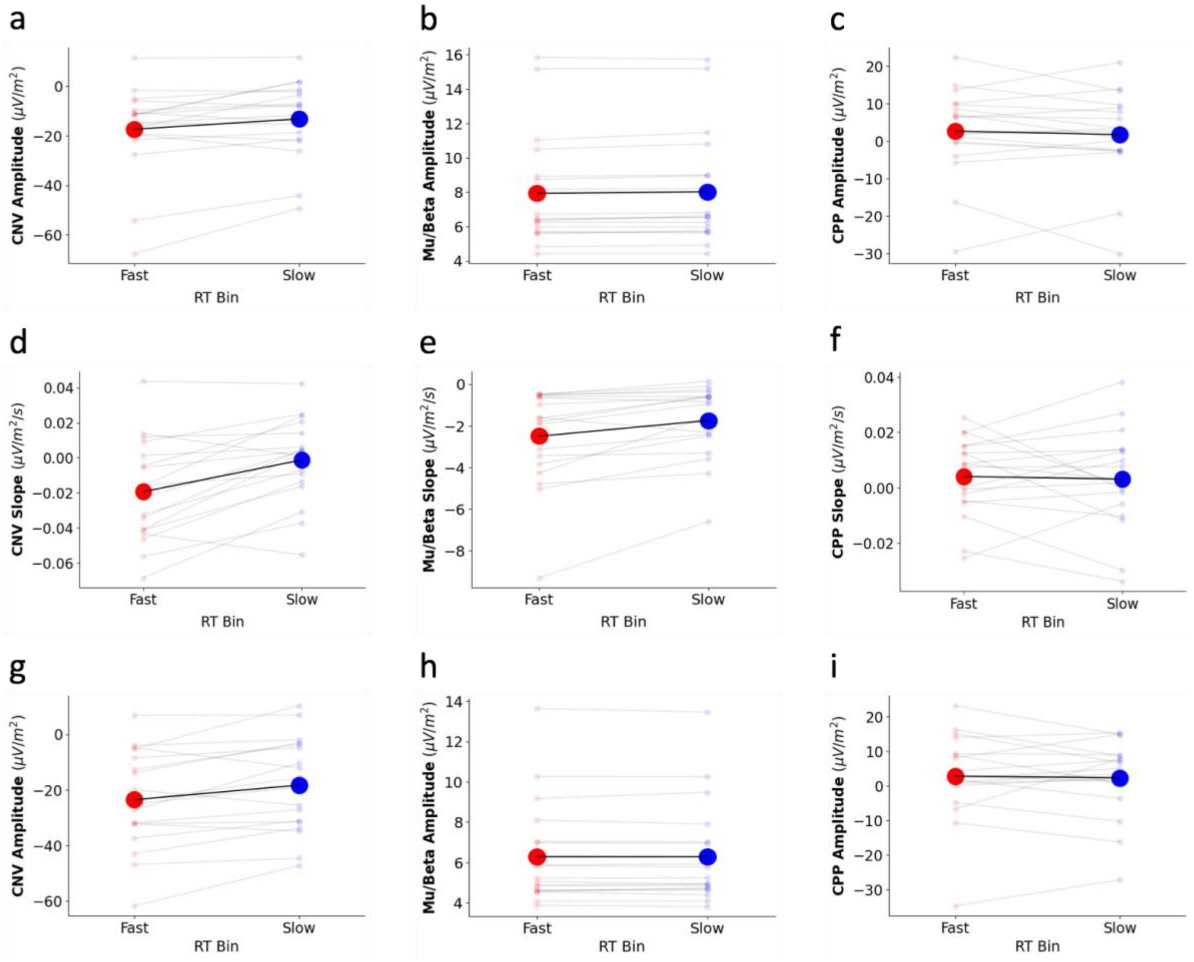


Figure S4: Group mean (black lines and large markers) and participant mean (grey lines and small markers) cue onset amplitude (first row), pre-response build-up rate (second row) and pre-response amplitude as a function of RT Bin. First column is CNV (a, d, g), second column is Mu/Beta (b, e, h) and third column is CPP (c, f, i).

#### S5: RT Bin ANOVAs

**Encode Offset**

- No effect of RT bin on CNV amplitude ($F(1, 17) = 1.80$, $p = 0.20$, $\eta^2_p = 0.1$)
- No effect of RT bin on CNV slope ($F(1, 17) = 3.7e^{-5}$, $p = 0.92$, $\eta^2_p = 5.48e^{-4}$)
- No effect of RT bin on Mu/Beta amplitude ($F(1, 17) = 4.28e^{-5}$, $p = 0.80$, $\eta^2_p = 0.004$)
- No effect of RT bin on Mu/Beta slope ($F(1, 17) = 1.66e^{-4}$, $p = 0.74$, $\eta^2_p = 0.006$)
- No effect of RT bin on CPP amplitude ($F(1, 17) = 0.97$, $p = 0.34$, $\eta^2_p = 0.05$)
- No effect of RT bin on CPP slope ($F(1, 17) = 1.8^{-5}$, $p = 0.95$, $\eta^2_p = 2.38e^{-4}$)

**Cue Onset**

- Significant effect of RT bin on CNV amplitude ($F(1, 17) = 6.43$, $p = 0.02$, $\eta^2_p = 0.27$)
- Significant effect of RT bin on Mu/Beta amplitude ($F(1, 17) = 7.86$, $p = 0.01$, $\eta^2_p = 0.32$)
- No effect of RT bin on CPP amplitude ($F(1, 17) = 0.46$, $p = 0.51$, $\eta^2_p = 0.03$)

**Reproduction**

- Significant effect of RT bin on CNV amplitude ($F(1, 17) = 10.50$, $p = 0.005$, $\eta^2_p = 0.38$)
- Significant effect of RT bin on CNV slope ($F(1, 17) = 19.09$, $p < 0.001$, $\eta^2_p = 0.53$)
- No effect of RT bin on Mu/Beta amplitude ($F(1, 17) = 3.85e^{-5}$, $p = 0.83$, $\eta^2_p = 0.003$)
- Significant effect of RT bin on Mu/Beta slope ($F(1, 17) = 12.74$, $p = 0.002$, $\eta^2_p = 0.43$)
- No effect of RT bin on CPP amplitude ($F(1, 17) = 3.69e^{-5}$, $p = 0.74$, $\eta^2_p = 8.42e^{-4}$)
- No effect of RT bin on CPP slope ($F(1, 17) = 4.34e^{-5}$, $p = 0.78$, $\eta^2_p = 0.005$)
